# Modulation of early host innate immune response by a Fowlpox virus (FWPV) lateral body protein

**DOI:** 10.1101/2020.10.02.324418

**Authors:** Efstathios S Giotis, Stephen M Laidlaw, Susanna R Bidgood, David Albrecht, Jemima J Burden, Rebecca C Robey, Jason Mercer, Michael A Skinner

**Affiliations:** Section of Virology, School of Medicine, St Mary’s Campus, Imperial College London W2 1PG, UK; School of Life Sciences, University of Essex, Colchester C04 3SQ, UK; Medical Research Council-Laboratory for Molecular Cell Biology, University College London, Gower Street, London WC1E 6BT, UK

**Keywords:** fowlpox virus, interferon, lateral bodies, nuclear localisation signal, microarrays

## Abstract

The avian pathogen, fowlpox virus (FWPV) has been successfully used as vaccine vector in poultry and humans but relatively little is known about its ability to modulate host antiviral immune responses in these hosts, which are replication permissive and non-permissive, respectively. FWPV is highly resistant to avian type I interferon (IFN) and able to completely block the host IFN-response. Microarray screening of host IFN-regulated gene expression in cells infected with 59 different, non-essential FWPV gene knock-out mutants revealed that FPV184 confers immunomodulatory capacity. We report that *FPV184*-knockout virus (FWPVΔ184) induces the cellular IFN response as early as 2 hours post-infection. The wild-type, uninduced phenotype can be rescued by transient expression of *FPV184* in FWPVΔ184-infected cells. Ectopic expression of *FPV184* inhibited polyI:C activation of the chicken IFN-β promoter and IFN-α activation of the chicken Mx promoter. Confocal and correlative super-resolution light and electron microscopy demonstrated that FPV184 has a functional nuclear localisation signal domain and is packaged in the lateral bodies of the virions. Taken together, these results provide a paradigm for a late poxvirus structural protein packaged in the lateral bodies and capable of supressing IFN induction early during the next round of infection.

## Introduction

Poxviruses are large, enveloped, double-stranded DNA viruses capable of causing disease in mammals, birds and insects. Binding and entry of poxviruses into vertebrate cells is an efficient process for a wide range of cell types, irrespective of the host species, with any host range restriction occurring after viral entry^1^. The complex replication cycles of poxviruses take place exclusively in the cytoplasm although it has long been suggested that poxviruses must interact with host nuclei for productive infection^2–6^. Perhaps the best-studied antiviral host restriction mechanism is interferon (IFN)-mediated, against which almost all viruses have evolved defence mechanisms^7,8^. Some of the first viral anti-IFN defence mechanisms were elucidated using VACV, which expresses multiple inhibitors of IFN induction, JAK/STAT signalling and IFN-stimulated genes (ISGs) as well as IFN-receptor antagonists and mimics of IFN ligands^7–12^.

These potent immunomodulators are produced mainly during the early phase of VACV gene expression. However, poxviruses have strategies in place to prevent or evade immediate early host innate responses induced as a consequence of the virus binding and fusing with the cell membrane. Poxviruses have proteinaceous sub-structures, termed lateral bodies (LBs), outside the core but within the mature virion’s membrane. These are analogous to herpesvirus tegument proteins, some of which perform immunomodulatory functions early during infection^13–16^.

Schmidt *et al.*^17^ reported that VACV packages the conserved H1 phosphatase (also known as VH1) within LBs. When VH1 is released from LBs into the cytoplasm of the host cell following membrane fusion, it acts to block IFN-γ mediated STAT-dependent signalling prior to gene expression^18^. Whether additional LB-resident viral immune modulators, capable of blocking other parts or the IFN system, are packaged in the LB of VACV or other poxviruses remained to be determined.

Relative to our understanding of the immunomodulation mediated by mammalian poxviruses, our knowledge of the strategies deployed by avipoxviruses to disarm the interferon response remains rudimentary. The prototypic member of the avipoxviruses, fowlpox virus (FWPV), is the causal agent of a widespread, enzootic disease of domestic chickens and other gallinaceous birds. It has been used as a recombinant vector for the expression of antigens from several avian and human pathogens^19^. In common with the other poxviruses, FWPV has developed strategies to disarm the host IFN-response and has been found to block efficiently the pI:C mediated induction of IFN-β promoter and the IFN-stimulated induction of ISGs in chicken cells^20^. Studies of FWPV immunomodulators have been complicated by the fact that only 110 (42%) of FWPV genes share significant similarity to those in other poxviruses^21^. To identify the innate immunomodulatory factors encoded by FWPV, we previously conducted two broad-scale pan-genome analyses of FP9, a highly attenuated strain used as a vaccine vector in both poultry and mammals^21,22^. In the first study, we identified *FPV012* as a modulator of IFN induction by screening a knock- out library of 65 non-essential FP9 genes^20,23^. In the second study, using a gain-of-function approach in which 4-8 kbp fragments of FP9 were introduced into modified vaccinia Ankara (MVA), we found that FPV014 contributes to increased resistance to exogenous recombinant chicken IFN-α^24^.

In this report, we used our existing FP9 knock-out library^20^ to screen infected primary chicken embryo fibroblasts (CEFs) for FWPV genes that modulate the induction of interferon-regulated genes (IRG)s. Using this approach, we identified FPV184 as a third FWPV immunomodulatory protein blocking the induction of innate immune responses. Intriguingly, unlike the FWPV immunomodulators FPV012 and FPV014 (which are both early viral proteins), FPV184 was found to be a late, structural protein with a functional nuclear localisation signal. Consistent with its ability to modulate ISG responses soon after infection and long before *de novo* production, we show that FPV184 is packaged into FWPV particles where it resides in the LBs. These results suggest that the packaging of late immunomodulatory proteins, and their subsequent delivery into the nucleus of newly infected cells serve as an immediate early innate immune evasion strategy.

## Results

### Identification of immunomodulatory signature induced by FWPVΔ184

We infected CEFs (three independent batches) with 59 individual FWPV mutants, each deficient in one non-essential gene^20,23^. Gene expression was analysed at 16 h post infection (p.i.) using Affymetrix Chicken 32K genome microarrays, which include probe sets for FWPV transcripts, allowing confirmation of viral infection and genotype for each knock-out virus. The parental FP9 strain blocks entirely the induction of IFN and ISGs and served as a negative control. The FPV012 knock-out (FWPVΔ012), which induces a subset of ISGs^20^, served as a positive control. All knock-out viruses were screened for their ability to induce ISGs, using a set of 337 chicken ISGs, which were determined by treating CEFs with IFNα for 6 h^25^ (ArrayExpress accession: E-MTAB-3711 and Supplementary Table S1).

FWPVΔ012 induced a subset of ISGs to moderate levels compared to the uninfected control (*n*= 87, FDR≤0.05, fold change≥1.5) and the FP9 virus (*n*= 98) (Fig. 1a-b). We found that FWPVΔ184 induced a smaller subset of ISGs compared with the uninfected control (*n*= 41) and the FP9 virus (*n*= 5; IFIT5, Mx1, IFI6, ARAHGAP8, LOC418700) (Fig. 1a-b and Supplementary Table S1).

**Fig. 1.**
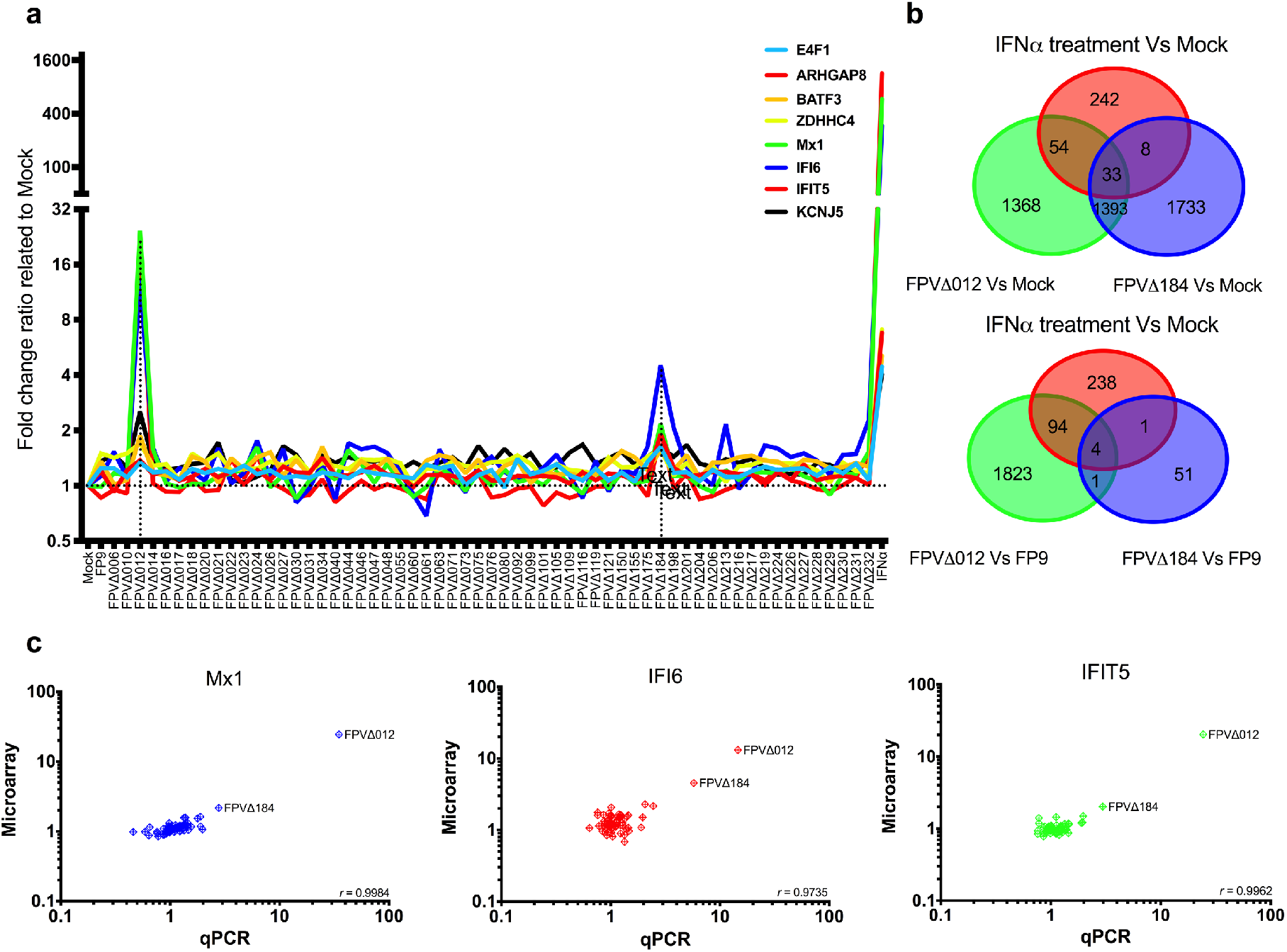
Disruption of *FPV012* (a known FP9 immunomodulator gene) and *FPV184* induced overlapping subsets of ISGs. **(a)** Gene expression of selected ISGs (E4F1, ARHGAP8, BATF3, ZDHHC4, Mx1, IFI6, IFIT5, KCNJ5Z) for all knock-out viruses measured by microarrays (fold-change relative to mock-treated CEF). **(b)** Venn diagrams, showing numbers of genes differentially regulated by FWPVΔ012, FWPVΔ184 or IFN, in comparison with mock-infected (upper) or parental FP9-infected (lower) cells. **(c)** Comparison of median differential expression levels for 3 selected transcripts determined by microarray and qPCR analysis (Mx1, IFI6, IFIT5). Plots show log10 expression fold change for the selected genes for CEF infected with all knock-out FP9 mutant viruses compared to mock-treated CEF. Pearson correlation coefficients (*r*) are shown in the lower right corner of each plot.

To confirm these results, we used real-time quantitative PCR (qPCR) to analyse mRNA levels of Mx1, IFI6 and IFIT5 upon infection with the knock-out viruses. Pearson’s correlation test (Fig. 1c) was performed to test for pairwise correlations between the two transcriptomic approaches. The correlation coefficients (*r*) for all comparisons were over 0.97, indicating the reproducibility of expression analysed by microarray or qPCR.

### *FPV184* is not an essential gene but its loss does impart a defect in viral growth

FPV184 is a small protein (88 amino acids) of predicted molecular weight 9.5 kDa. The presence of a late promoter, TAAATG, upstream of *FPV184* suggested that it is a late-expressed gene unlike the other two known FWPV immunomodulators *FPV012* and *FPV014*, which are expressed early during viral replication. Microarrays showed that *FPV184* is indeed expressed at intermediate/late stages of FWPV replication (Supplementary Fig. S2c). To determine if *FPV184* is essential for FWPV replication, we used two recombinant FWPV viruses containing a disrupted *FPV184* gene, a PCR-mediated knock-out virus (FWPVΔ184, used in microarrays and in the rest of the study) and a transient dominant virus (TDdel184). These deletion viruses could be isolated in culture, indicating that *FPV184* is non-essential for growth *in vitro* and both are able to induce Mx1 in CEF (Fig. 2a). However, the viruses had lower growth rates (approx. 0.75 log10 lower), reduced viral yield in CEFs (approx. 1-1.5 log10 lower) and smaller plaque diameters (approx. two-thirds) compared to the parental FP9 virus (Fig. 2b-e) indicating that loss of *FPV184* imparts a growth defect on FWPV.

**Fig. 2.**
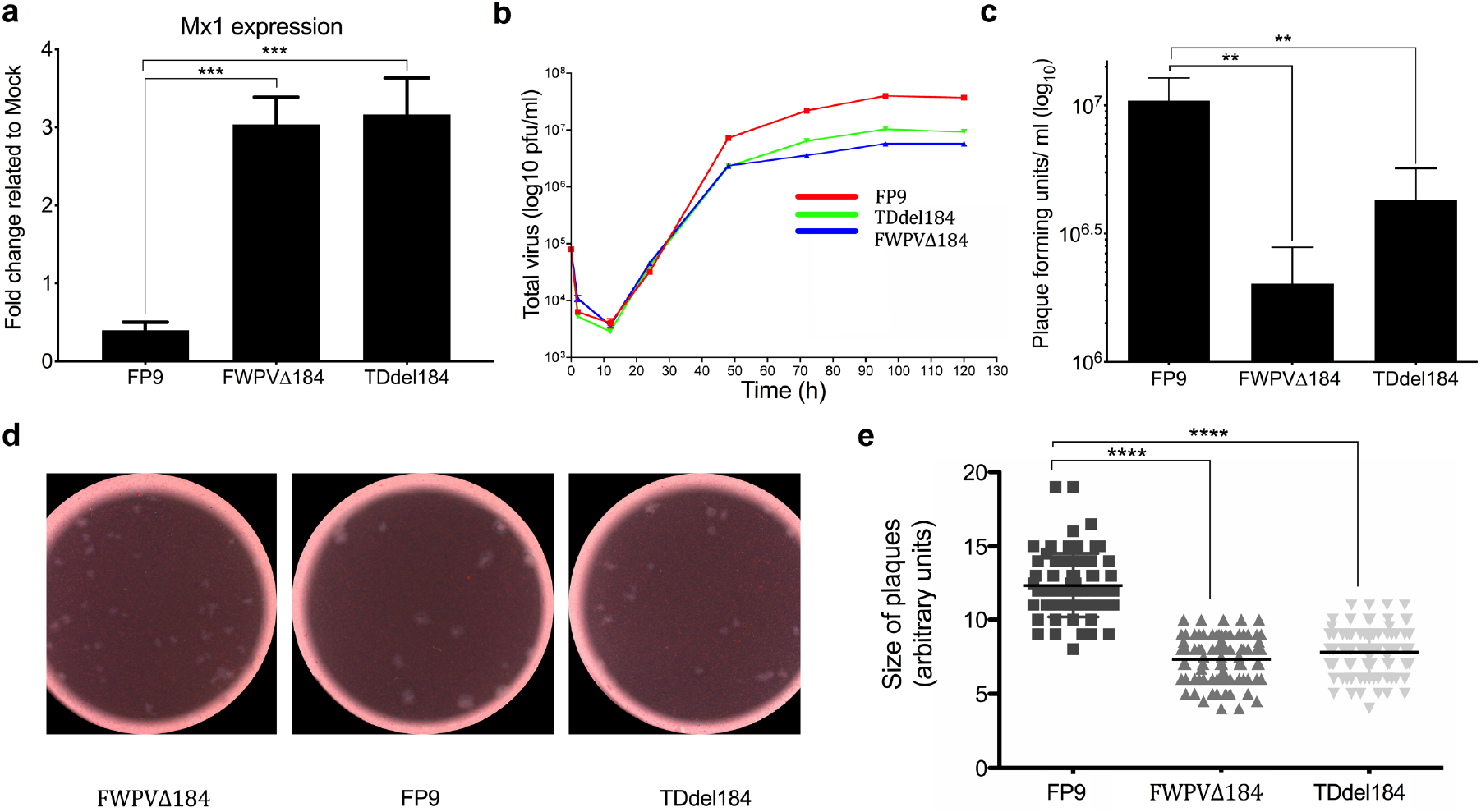
Deletion of *FPV184* affects the replicative fitness of the recombinant virus. **(a)** Independent transient dominant (TDdel184) and GPT-insertion *FPV184* knock-out mutant (FWPVΔ184) viruses (MOI: 5, 1 h adsorption) induce similar mRNA levels of Mx1 at 4 h p.i. in CEF compared to parental FP9 virus by qPCR. **(b)** Multiple-step growth analysis of FP9, FWPVΔ184 and TDdel184 viruses in CEF (MOI: 0.01). **(c)** Viral yield of FP9, FWPVΔ184 and TDdel184 viruses (in pfu/ml) following 72 h infection with an MOI: 0.1 in CEF. **(d)** Images of viral plaques of FP9, FWPVΔ184 and TDdel184 viruses in CEF and **(e)** Scatterplot showing their respective size in arbitrary units. **(a-e)** Error bars indicate Standard Error from the Mean (SEM); *n*=3. One-way ANOVA with Dunnett’s *post hoc* test were used to analyse the data. \*\**p*< 0.01, \*\*\**p*< 0.001, \*\*\*\**p*< 0.0001.

### FPV184 mediates early suppression of ISGs through LB packaging

The expression kinetics of ISGs in CEFs infected with FWPVΔ184 or FP9 were monitored for 16 h p.i. using qPCR. FP9 infection reduced basal Mx1 expression by 30% throughout the course of infection (Fig. 3a shows mRNA expression of Mx1 in infected with FP9 and FWPVΔ184 related to uninfected samples). Conversely, cells infected with FWPVΔ184 showed a modest bimodal increase in Mx1 expression from 2 to 5 h p.i. (2.8-fold compared with uninfected and 4.2-fold compared to FP9), and again at 14 to 16 h p.i. (2.2-fold compared with uninfected and 3.1-fold compared to FP9; Fig. 3a). Immunoblot analysis at 4 and 16 h p.i. confirmed that FWPVΔ184 could not suppress Mx1 protein expression (Fig. 3b). As FPV184 is only expressed later during infection (Fig. 3c and Supplementary Fig. S2) we confirmed late protein expression by immunoblot against another late protein (FPV191; Fig. 3b), which confirmed that no late viral protein was expressed at 4 h p.i.

**Fig. 3.**
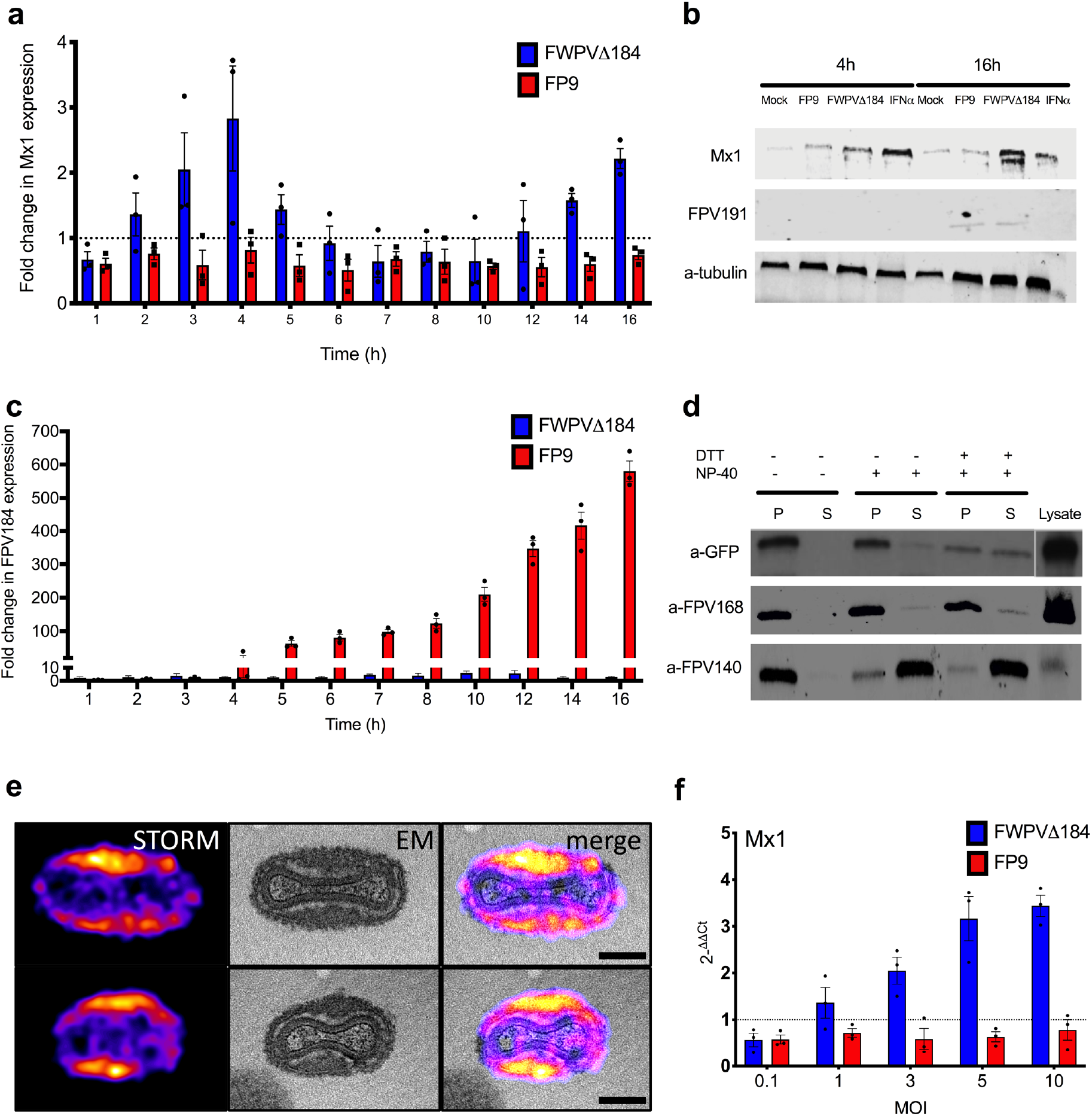
FPV184 mediates early suppression of ISGs through LB packaging. **(a)** Kinetics of Mx1 mRNA expression (means ± SEM, *n*=3) following infection of CEF with FWPVΔ184 and the wild-type FP9 viruses (MOI: 5, 1 h adsorption), measured by qPCR. **(b)** Western-blot showing protein expression of the chicken Mx1, the late-expressed FPV191 and α-tubulin at 4 and 16 h p.i. infection of CEF with FP9 and FWPVΔ184 (MOI: 5) and treatment with IFN-α (1000 IU/ml). **(c)** Kinetics of expression of FPV184 expression (means ± SEM, *n*=3) following infection of CEF with FWPVΔ184 and the wild-type FP9 viruses measured by qRT-PCR. **(d)** Purified GFP184 virions were treated with either NP-40 and/or DTT, centrifuged to separate the pellet (P) from supernatant (S) fractions, then subjected to SDS/PAGE and western blotting. Immunodetection was performed with α-GFP antiserum to detect GFP fusion protein(s) and monoclonal antibodies against FWPV structural proteins FPV140 (localised on mature virions’ membrane) and FPV168 (localised in the virion core)^27^. **(e)** Correlative super-resolution light and electron microscopy of GFP184 virions immunolabelled with anti-GFP nanobodies. STORM images of the lateral body protein were registered with EM micrographs. Two representative virions, are shown at higher magnification in upper and lower panels (i and ii). Scale bars: 300 nm. Overview is shown in Supplementary Fig. S3. **(f)** mRNA expression levels of Mx1 in CEF at 4 h p.i. infected with FP9 or FWPVΔ184 using different MOI (0.1, 1, 3, 5 and 10), measured by qPCR. Columns represent fold-change (means ± SEM, *n*=3) compared to mock-treated CEF.

This late expression of FPV184 is unusual, if not currently unique, for a non-essential poxviral gene with early immunomodulatory function. The only known example of an immunomodulatory poxviral protein with an early effect is the essential VACV H1 phosphatase, which is packaged into poxvirus LBs and delivered into host cells during virus entry to mediate early suppression of STAT1 signalling^17^. Thus, we asked if FPV184 is packaged into FWPV virions and, if so, where in the virions it is located. Purified GFP184 virions (expressing FPV184 with GFP fused to its N-terminus) were left untreated or were subjected to fractionation using NP-40 and/or DTT to separate viral membranes from cores and their associated LBs^26^. Immunoblots directed against FWPV core protein FPV168, and membrane protein FPV140 were used to validate the fractionation^27^. Immunoblots directed against GFP indicated that GFP184 is packaged in virions (Fig. 3d). In untreated and NP-40-treated virions, very little GFP184 was released from virions, suggesting that the protein is not on the virion surface. Treatment with NP-40 and DTT^28^ resulted in a 50/50 distribution of GFP184 between virion membrane and core/LB fractions, suggesting that GFP184 resides between the viral membrane and the viral core.

To more accurately define the sub-viral localisation of FPV184 we used correlative super-resolution light and electron microscopy (CSRLEM). Purified GFP184 virions were imaged using stochastic optical reconstruction microscopy (STORM) followed by transmission electron microscopy (TEM). Images were registered to correlate the fluorescence signal with EM structural information (Fig. 3e; Supplementary Fig. S3). STORM images showed that GFP184 was localized to the two distinct LB structures running the length of the virions and absent from the virus cores. Correlation of these images with the corresponding TEM images showed that GFP184 is a LB resident protein. Consistent with its delivery via LBs, the activation of Mx1 gene expression at 4 h p.i. in cells infected with virions lacking FPV184, was found to be dose (MOI)-dependent (Fig. 3f).

### Ectopic expression of FPV184 inhibits polyI:C- and IFN-stimulated activation of chicken IFN-β and Mx promoters, respectively

To directly assess the role of FPV184 in the absence of other immunomodulatory proteins expressed during infection, we used a construct (pFPV184) to overexpress FPV184 in immortalised DF-1 chicken fibroblast cells and assessed its ability to modulate the induction of the chicken IFN-β promoter by polyI:C (Fig. 4a) or the chicken Mx promoter by recombinant IFN-α, as detected using a luciferase reporter assay (Fig. 4b). Compared to the empty vector, pFPV184 inhibited induction of the chicken IFN-β promoter and the Mx promoter by 44% and 22%, respectively. Overexpression of FPV012, which was used as a positive control for these experiments inhibited induction by 74% and 51% for IFN-β and Mx promoters, respectively. Confirming its role in ISG suppression during infection, transient expression of FPV184 in DF-1 cells infected with the FWPVΔ184 resulted in wild type ISG-suppression levels (Fig. 4c). Co-transfection of the immortalised DF-1 chicken fibroblast cells with constructs expressing FPV184 and/or the other known FWPV immunomodulators, FPV012 and FPV014, showed that there is no synergism between the three proteins in inhibiting the IFN-α stimulation of Mx1 promoter (Supplementary Fig. S4).

**Fig. 4.**
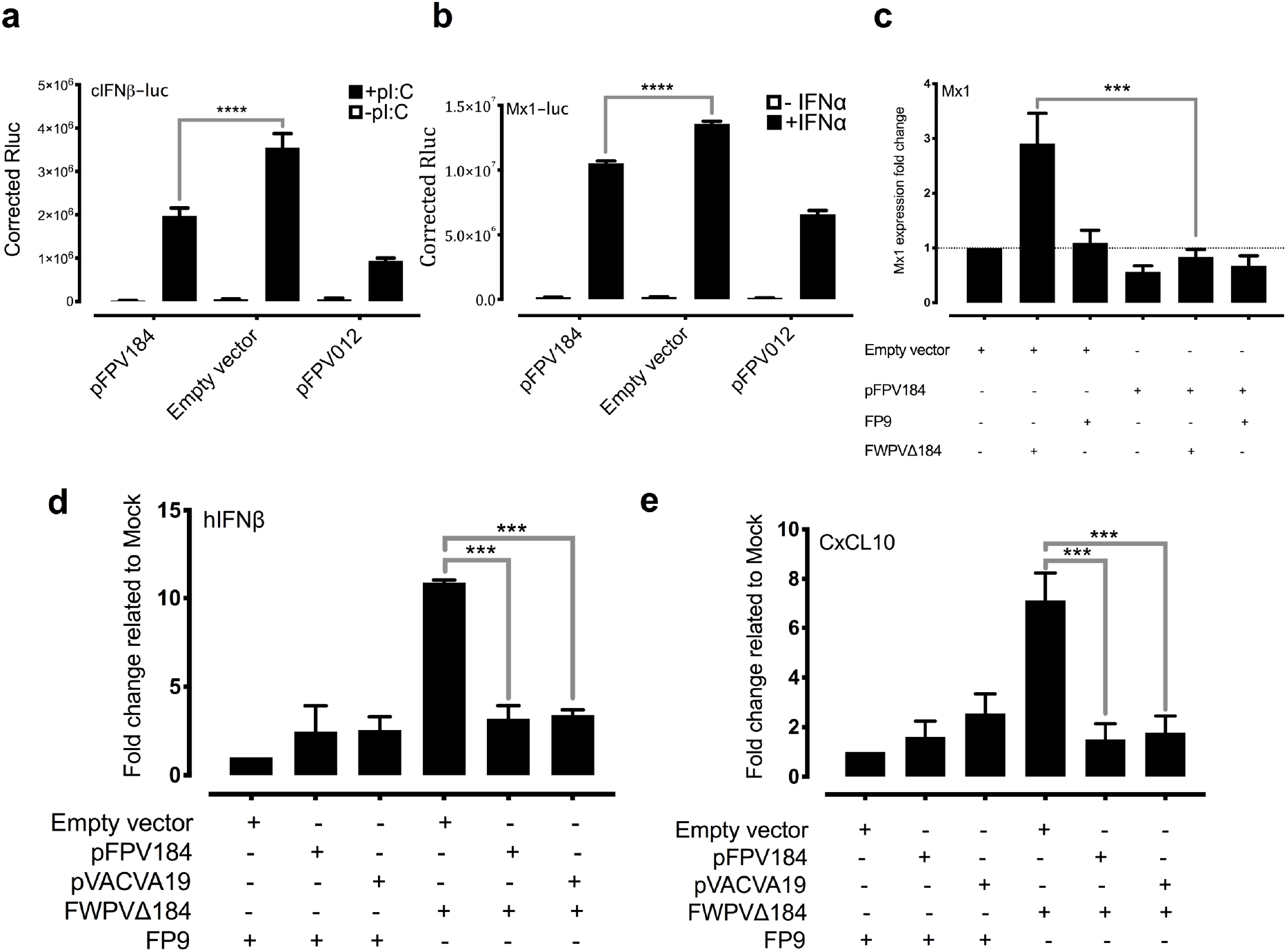
Ectopic expression of FPV184 elicits an immunomodulatory effect in chicken and human cells. Using a luciferase assay, the FLAG-tagged FPV184 protein was shown to inhibit induction of both IFNβ promoter by polyI:C **(a),** and to a lesser extent the Mx1 promoter by IFNα **(b)** in chicken DF-1 cells. Two-way ANOVA (type of plasmid and effect of IFNα or pI:C) with Bonferroni *post hoc* test were used to analyse the data (**** *p*<0.0001). **(c)** Complementation assay measuring Mx1 induction by qRT-PCR in DF-1 cells transiently transfected (48 h) with either the FPV184 expression plasmid or with the empty vector and infected with FWPVΔ184 or FP9 for 4 h (MOI: 5). Results shown are representative of three independent experiments. One-way ANOVA with Tukey’s *post hoc* test were used to analyse the data (\*\*\**p*<0.001). **(d)** qPCR analysis of human IFN-β, and **(e)** CXCL10 mRNA levels in HEK293T cells transfected for 48 h with either an expression plasmid for FPV184 or VACV A19 and infected with parental FP9 or FWPVΔ184 for 4 h (MOI: 5). Error bars indicate SEM; *n*=3. One-way ANOVA with Dunnett’s *post hoc* test were used to analyse the data. \*\*\**p*<0.001.

### Ectopic expression of FPV184 and its ortholog, VACV A19, restores the immunomodulatory capacity of FWPVΔ184 in human cells

FPV184 is a well-conserved orthologue of VACV A19 (~ 40% amino acid identity, Supplementary Fig. S5), an essential, late structural protein found in vertebrate but not insect poxviruses^29,30^. To ask if VACV A19 could substitute for FPV184, VACV A19 was overexpressed in human HEK293T cells infected with FWPVΔ184 (Fig. 4d-e; Supplementary Fig. S6). Under these conditions, expression of VACV A19 suppressed the induction of human IFN-β and CXCL10 as efficiently as FPV184, which served as a positive control for this experiment. The ability of A19 to complement for the loss of FPV184 suggests that these proteins act as functional equivalents to suppress early immune responses upon infection.

### FPV184 contains a functional NLS partially responsible for its immunomodulatory activity

It has been previously reported that VACV A19 and its orthologs, including FPV184, contain 3 highly conserved motifs: Two CxxC motifs in the middle of the protein (amino acids 37-40 and 72-75), whose mutation resulted in a reduction in virus yield, and a basic NLS (FWPV amino acids 9-14: KKRKKR; Fig. 5a and Supplementary Fig. S5) at its N-terminus^30^. While A19 was shown to display nucleo-cytoplasmic localization during infection, mutation of the NLS had no apparent defect on virus growth^30^.

**Fig. 5.**
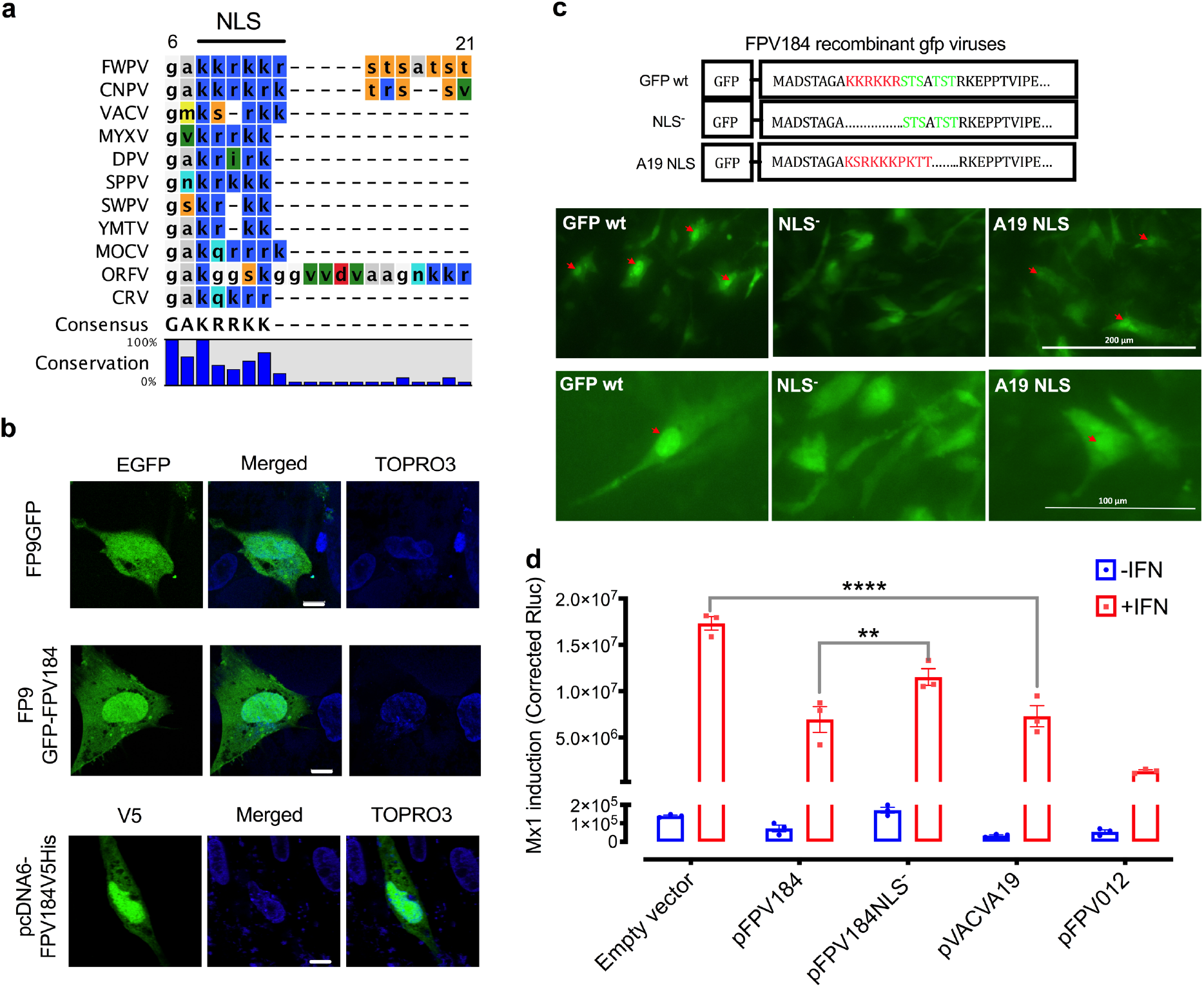
The FPV184 NLS is functional and nuclear localisation is needed for the protein’s immunomodulatory ability. **(a)** Multiple amino acid sequence alignment of FPV184 orthologues illustrated an NLS motif. Detailed information on sequences and full alignment can be found in the materials and methods and Supplementary Fig. S5.**(b)** Representative confocal microscopy images of CEF transfected for 48 h with expression plasmids containing either a GFP tag alone (FP9GFP), or a GFP-tagged FPV184 (FP9GFP-FPV184) and infected with FP9 for 16 h or transfected for 48 h with expression plasmid containing V5-tagged FPV184 (pcDNA6FPV184V5His) but left uninfected. Scale bar, 8 μm. **(c)** Upper panel: Schematic depicting modifications of the FPV184 N-terminal region in the recombinant GFP viruses (NLS is shown in red; the putative phosphorylation sites in green). Lower panel: Widefield fluorescence micrographs of CEF 24 h p.i. at a MOI of 1 with the recombinant GFP viruses. Red arrows show nuclear-localised fluorescence. **(d)** Luciferase signals in DF-1 cells after transfection with chicken Mx1-luciferase reporter plasmid together with expression plasmids for FPV184, FPV012, VACVA19 and FPV184 without the NLS domain (pFPV184NLS^−^). Two-way ANOVA with Tukey’s *post hoc* test were used to analyse the data (\*\**P* < 0.01, \*\*\*\**P* < 0.0001). Error bars indicate SEM; *n*=3.

To study the cellular localisation of FPV184 during infection, a construct expressing GFP/cMyc-tagged FPV184 under the control of an early/late synthetic poxvirus promoter was generated^31^. Consistent with the VACV A19 data, expression of this tagged version of FPV184 in FP9-infected CEFs showed its presence throughout the cytoplasm but concentrated within the nucleus (Fig. 5b). A non-fused EGFP control, was fairly evenly distributed throughout the cytoplasm and nucleus. Expression of V5-His-tagged FPV184 from a eukaryotic expression vector, in the absence of infection, showed the same distribution (Fig. 5b).

To determine whether the identified FPV184 NLS motif was responsible for this nuclear accumulation, we transiently expressed FPV184 lacking its NLS (Fig. 5c; NLS^−^). To determine if the nuclear localisation signal found in FPV184 is functionally conserved with VACV A19, we also swapped this region of FPV184 with the corresponding sequence from VACV WR A19 (KSRKKKPKTT) (Fig. 5c; A19 NLS). These constructs were used to generate FWPV recombinants expressing the GFP-tagged versions of FWPV184 as second copies. Following infection with these viruses, GFP184 showed diffuse cytoplasmic staining and distinct accumulation within both the viral factory and nucleus, while expression of the NLS^−^ recombinant GFP viruses produced a diffuse fluorescence within infected cells (Fig. 5c & confocal microscope analysis in Supplementary Fig. S7). Infection of cells with A19 NLS resulted in nuclear localisation, although the intensity was decreased compared to parental GFP184 (Fig. 5c & Supplementary Fig. S7). These results suggest that FWPV184 contains a functional NLS that is responsible for localizing a steady state portion of the viral protein into the cell nucleus throughout infection.

To assess the impact of the unusual FPV184 nuclear localization on its ability to suppress Mx1, FPV184, FPV184 NLS- and FPV184 A19 NLS were transiently expressed in DF-1 cells and a Mx1-luciferase reporter assay performed in the absence or presence of recombinant chicken IFNα (Fig. 5d). FPV184 partially suppressed IFN-mediated induction of the chicken Mx1 promoter, while FPV184 NLS- was reduced in its ability to suppress the promoter (Fig. 5d). Consistent with its conserved sequence, VACV A19 NLS was able to rescue suppression by FPV184 A19 NLS to the level observed with FPV184 (Fig. 5d).

## Discussion

A common strategy to generate new or improved live recombinant poxvirus vaccines is to target and delete immunomodulatory genes in the vector. There has been considerable study of multiple immunomodulatory proteins of VACV^8–10,12^ but, in contrast, few immunomodulators have been reported in other poxviruses. In FWPV, two immunomodulatory proteins have been reported so far: FPV012 and FPV014^20,24^. Both are members of the ankyrin repeat protein superfamily (Pfam clan CLO465), are expressed early during viral replication and are not essential for virus replication in culture.

In a systematic, large-scale study we interrogated the innate immune function of non-essential FWPV genes, by transcriptomic analysis of a library of KO viruses derived from the highly attenuated FWPV vaccine strain FP9. We identified a third immunomodulatory protein (FPV184), which, unlike the other two immunomodulators, is not an ankyrin protein, is expressed late during FP9 replication and encodes a very small structural protein packaged in the virions. There is a good degree of conservation between FPV184 and its VACV orthologue A19. A19 is also expressed at the intermediate/late stages of replication but, unlike FPV184, it has been reported as essential for viral replication^30^, being involved in the maturation of VACV virions and in early viral transcription in newly-infected cells, where it interacts directly with the viral RNA polymerase and other members of the early transcription complex^29,30^. Since A19 is a late protein and has an essential role in virion morphogenesis, A19 knock-out viruses were not constructed and thus the potential role of A19 as an immunomodulator was not evaluated. Whilst conservation of A19 and FPV184 suggests an important function for the latter, we were able to generate FWPV lacking *FPV184* using two different methodologies, showing that the gene is non-essential for replication *in vitro*, albeit with a reduced replication rate.

Like VACV A19, FPV184 may act as an early viral transcription factor or a regulator. The two CxxC motifs involved in binding of the VACV orthologue to viral RNA polymerase^29,30^ are well conserved in FPV184. Although not investigated in this study, the presence of CxxC motifs is indicative of a zinc finger motif^32,33^. The zinc-binding domains, initially identified as DNA binding motifs in transcription factors, are now grouped into super-families based on their amino-acid composition. A number of transcription factors have been found to bind the DNA sequence (A/T)GATA(A/G) in the regulatory region of genes. These have been termed GATA-binding transcription factors and are able to bind DNA *via* a conserved zinc-finger domain in which the zinc ion is coordinated by 4 cysteine residues^34^. In VACV, early transcription factor null mutants have displayed defects in morphogenesis^30,35,36^. We have not conducted morphological studies of the FWPVΔ184 virions, *e.g.* by using electron microscopy, consequently a role for FPV184 in morphogenesis cannot be excluded.

It is well established that FWPV blocks entirely the launch of avian type I IFN response and the induction of ISGs. Therefore, the relatively subtle induction of a subset of ISGs by FWPVΔ184 could be explained by the presence of a multigenic, redundant system in FP9 to control host interferon response, as observed in VACV and other poxviruses, or by relatively mild stimuli. It is possible that the complementary effects of other FWPV immunomodulatory proteins alleviate the effects caused by the absence of FPV184. For example, FPV138 is a homologue of VACV H1, a protein-tyrosine phosphatase that resides in the LBs, which, upon release into the cytoplasm, dampens type I IFN-induced, STAT1-mediated signalling. ISG expression was higher in cells infected with FWPVΔ012 than with FPVΔ184, in terms of both numbers and expression levels; there was, however, a significant overlap between the ISGs induced by both viruses (Fig. 1b). Furthermore, when FPV012 and FPV184 were transiently co-expressed in DF-1 cells, they were not synergistic in inhibiting the pI:C-mediated induction of IFN-β (Supplementary Fig. S4), suggesting they may target the same host immune pathways but at different time points during infection. FPV184’s role may be to shut down immediate-early host immune responses and its function may be gradually superseded by that of the early immunomodulatory proteins expressed *de novo* upon infection.

The putative role of FPV184 in blocking immediate-early host immune response is also supported by our demonstration, with super resolution microscopy and CSRLEM, that the protein is packaged within the LB and outside the confines of the viral cores where early viral transcription is executed. LB have been described as a poxvirus mechanism for delivery of viral proteins to the cytoplasm of the cell soon after fusion of the MV membrane with cellular plasma or endosomal membranes; their release and disaggregation is believed to depend upon reduction and proteosomal activity^17^. Parallels exist with the herpes virus tegument, which is also located inside the virion, under the envelope but outside of the capsid, and is known to deliver virion host effector proteins into cells^17,37^. Our hypothesis is that FWPV LB contain additional packaged immunomodulatory factors that can act before core activation and early gene expression, to establish a favourable environment in the cytoplasm. Other proteins found in LB in VACV^17^ include the phosphoprotein F17 and oxidoreductase G4, both involved in morphogenesis, as well as the dual-specificity phosphatase H1 discussed above (FWPV orthologues, FPV103, FPV077 and FPV138, respectively). H1 has immunomodulatory function by virtue of its ability to target STAT-mediated signalling, required for IFN-mediated induction of ISG expression. Our demonstration that FPV184 and VACV A19 are also packaged in LB, with functional evidence that they both have immunomodulatory function, shows that they could complement H1, primarily by blocking induction of IFN but, to a lesser extent, also by blocking IFN-mediated induction of ISGs.

Another unexpected observation was that FPV184 is preferentially localised in the nuclei of host cells. Few host nuclear proteins have been shown to play a part in the poxvirus lifecycle and fewer poxviral proteins (*e.g.* VACV C6, VACV F16) have been found to enter the nucleus during an infection^38,39^. There is no evidence that host proteins are required either for DNA replication or early gene transcription^40^, but it is known that host proteins are necessary for VACV post-replication transcription; intermediate transcription requires host protein VITF-2, which resides within the nucleus of uninfected cells^41^. Although some poxviral genes have been reported to enter the nucleus, to date there has been no identification of a poxvirus-encoded protein containing an identifiable and functional NLS. Furthermore, the nuclear localisation of FPV184 was found to influence its immunomodulatory ability. Using a luciferase assay, we showed that a construct expressing FPV184 without the NLS only partially abrogated the ability of the protein to inhibit the IFN-mediated induction of Mx1 promoter (Fig. 5d). Whether nuclear localization is partially or fully required for the immunomodulatory ability of FPV184 was unclear as, due to its small size (9.5 kDa), low levels of the protein can enter the nucleus without an NLS, *via* passive diffusion. Myxoma virus encodes a protein termed myxoma nuclear factor (MNF)^42^, an ankyrin repeat containing protein that localises to the nucleus in the absence of an NLS and sequesters NFκB. The cowpox virus protein crmA, even though it is small enough to shuttle between the nucleus and cytoplasm by passive diffusion, requires a leucine-rich nuclear export signal (NES) for its nuclear export^43^. It is possible that, in the presence of the active NLS, the accumulation of FPV184 in the nucleus is dependent on its lack of a NES.

Although the conserved block of lysines and arginines found at the N terminus of FPV184 is characteristic of an NLS, it could also constitute a DNA/RNA binding domain. The possibility that FPV184 might be a dsRNA-binding protein could explain the inhibition of polyI:C mediated-induction of the IFN-β promoter but would not explain the inhibition of IFN-mediated induction of the Mx1 promoter we observed.

Collectively, the findings of this study indicate a late-expressed poxviral protein, packaged outside of the viral cores, with a non-essential phenotype, a functional NLS and an immediate early effect on the innate immune response; properties that make FPV184 resemble a herpesvirus tegument protein rather than a typical early, immunomodulatory poxviral protein. The precise mechanism(s) and function(s) of FPV184 are obscure. Nevertheless, these findings extend the paradigm by which poxvirus structural proteins can block induction of innate immune responses, to immediately after infection, and might be useful in vaccine development.

## Methods

### Cells and viruses

Freshly isolated chicken embryo fibroblasts (CEF)s were provided by the former Institute for Animal Health, Compton, Berks, UK (now the Pirbright Institute, UK). CEF were cultured in 199 media (Gibco^®^, Invitrogen) supplemented with 8% heat-inactivated newborn calf serum (NBCS; Gibco®, Invitrogen), 10% tryptose phosphate broth (Sigma), 2% nystatin (Sigma) and 0.1% penicillin-streptomycin (Gibco^®^, Invitrogen). DF-1 and HEK293T cells were maintained in 10% fetal bovine serum (Gibco^®^, Invitrogen), and 0.1% penicillin-streptomycin. All cells were grown at 37°C. The origin, propagation and titration (plaque assays) of FP9 virus have been described previously^21,44,45^. All transient transfections in the study were conducted with Fugene HD Transfection reagent (Promega) unless otherwise specified. To collect purified virus, confluent CEF were infected with FP9 at 0.1 PFU per cell. At 5 days p.i., the supernatant was harvested along with remaining cells. Virus purification has been described previously^45^.

### Multi-step growth curve

Confluent CEF were infected with the PCR mediated (FWPVΔ184) and transient dominant (TDdel184) knock-out viruses of FPV184 or with the wild-type virus (FP9) at 0.01 PFU per cell. The inoculum was removed 1 h later and replaced by fresh medium. At different times p.i., the extracellular medium was collected, and the cells were overlaid with 1 ml of fresh medium and stored at −70°C. Intracellular and extracellular viruses were subjected to titer determination by plaque assay^46^. Plaque sizes between wild-type and knock-out viruses were evaluated as before. Briefly, plaques areas were digitally enlarged and calculated in arbitrary units using ImageJ v.32 image analysis software (Fig. 2d). Scatterplots were created with GraphPad prism (Fig. 2e).

### Construction of plasmids and recombinant viruses

Construction of *FPV184* knock-out virus was done using two different approaches: a PCR mediated knock-out with a guanine-phosphoribosyltransferase (GPT) insertion in the middle of the gene (FWPVΔ184, which was used for the microarray study) and a transient dominant deletion knock-out virus (TDdel184)^47^, to produce recombinant viruses respectively. The resulting knockout viruses were sequenced in the region of *FPV184*, paying attention to the overlap with *FPV183*, to check that no adventitious mutations had been introduced inadvertently during the knock-out procedure. The nearest flanking genes to *FPV184*, *FPV183* and *FPV186*, are oriented such that the termini of the genes are towards *FPV184*, therefore making it unlikely that manipulation of *FPV184* would disrupt promoters of any flanking genes. Construction of dually tagged *FPV184* was not possible as the sequence encoding the C-termini of *FPV183* and *FPV184* overlap by [about] 30 bp. In confirmation of the screen, both recombinant viruses lacking FPV184 could no longer suppress Mx1 relative to FP9 virus infection (4 h p.i.; Fig. 2a).

#### Isolation of FPV184 deletion mutants by PCR-mediated knock-out

All primers used in the study are listed in the Supplementary Table S8. Insertional mutagenesis has been described previously^20,23^. PCR-mediated knock-out of *FPV184* was carried out using three sets of primers; FWPV DNA was used as the template for PCR with primers M2840 to M2841 and M2844 to M2845 whilst pGPTNEB193rev was used as the template for primers M2842 to M2843. The resultant PCR products were combined in equimolar amounts and a further round of PCR was carried out using the flanking primer pairs M2840 to M2845. The full-length PCR construct was then used to transfect FP9 infected-CEF and the recombinant viruses were selected using media containing mycophenolic acid (25 μg ml^−1^), xanthine (250 μg ml^−1^), and hypoxanthine (15 μg ml^−1^) (MXH). Virus was passaged in T25 flasks three times in the presence of MXH before plaquing. Passaged virus was then plaque purified once in the absence of MXH and once more in the presence of MXH.

#### Isolation of FPV184 deletion mutants by transient dominant selection

The *FPV184* deletion plasmid was constructed using the previously described vector pGNR^44^. The 5’ end of the *FPV184* gene and 200 bp of upstream sequence was amplified by PCR using primer pairs M2854 to M2856 and the 3’ end of the *FPV184* gene and 200 bp of downstream sequence was amplified using primer pairs M2857 to M2855. Following this first round of PCR the products were combined in equimolar amounts and a second round of PCR carried out using the flanking primer pair M2854 to M2855. Utilising *BamH*I and *Hind*III sites within M2854 and M2855 respectively the second round PCR product was digested and cloned into pGNR/ *BamH*I/ *Hind*III to produce pGNRFPVΔ184. Deletion mutants were isolated by the transient dominant selection method^47^ as described previously^44^.

#### Confirmation of FPV184 deletion mutants

Each deletion mutant was screened by PCR with flanking primers (giving PCR products of specific sizes for wild-type and deleted genes), one flanking primer and one primer internal to the deletion (detecting only the wild type gene) or one flanking primer and one primer specific for the *GPT* gene (detecting insertion of *GPT* into the wild type gene). The primers used are as follows: flanking primers M530 TO M1257, internal primer M2919 to M1257, PCR mediated knock-out *GPT* primers M192 to M2854, transient dominant knock-out *GPT* primers M192 to M1257.

#### Generation of GFP recombinant viruses

The *FPV184* gene was amplified by PCR with M2952 and M2951. The product was digested with *Xma*I and *Sac*II (within M2952 and M2951 respectively) and cloned into pEFGPT12S-CvectorEGFPmyc/*Xma*I/*Sac*II to produce pCVecGFP184 (GFP at the N-terminus). Mutations were introduced into the *FPV184* gene by PCR using the following primer pairs (as shown in Fig. 5c): M2953 and M2951 for NLS^−^; M2956 and M2951 for A19 NLS and two more controls to assess the effect of a putative phosphorylation site (PT; results not shown; M2954 and M2951 for NLS^−^/PT^−^; M2955 and M2951 for PT^−^. All of the products were digested with *Xma*I and *Sac*II and cloned into pEFGPT12S-CvectorEGFPmyc/*Xma*I/*Sac*II to produce the mutant viruses. Following transfection of constructs into FP9-infected CEF, recombinant viruses were selected for, using mycophenolic acid, and plaque purified twice.

#### Cloning for ectopic expression of viral genes

The *FPV184* gene was amplified by PCR with M2892 and M2893. The product was digested with *Hind*III and *Xho*I and cloned into pcDNA6V5His/*Hind*III/*Xho*I (Invitrogen) to give pcDNA6FPV184V5His or amplified with 4279 and 4280 and digested with *XhoI* and *SalI* into pCIFLAG(N-terminus)/*XhoI*/*SalI* to give pFPV184. VACV A19 was amplified with 4344 and 4345, digested with *XhoI* and *SalI* into pCIFLAG/*XhoI*/*SalI* to give pVACVA19.

### Infection of CEF for microarray and qPCR analyses

Media was removed from fully confluent CEF (in T25 flasks; Greiner Bio One; 5.6 × 10^6^ cells/flask) and replaced with 8 ml DMEM containing DMEM (Mock), FP9 (MOI: 5) or one of the knock-out viruses (MOI: 5). After 2 h, DMEM or virus-containing DMEM was replaced with culture media (199 media supplemented with 2% NBCS, 10% TPB, 2% nystatin and 0.1% penicillin/streptomycin) and cells were then incubated for a further 14 h before harvesting. Mock- and virus-infected cells were harvested at 16 h post infection and stored at −80°C in RNALater (Sigma) until RNA extraction. The experiment was repeated in triplicate for each knock-out virus (for FWPVΔ228 virus duplicates were used) using three different batches of CEF.

### RNA extraction and processing of samples for microarray

RNA isolation and processing of samples and microarrays was done as previously described^48^. Total RNA was extracted from mock-, infected-, and IFN-stimulated DF-1 and CEF using the RNeasy kit (Qiagen) according to the manufacturer’s instructions as previously described^48^. On-column DNA digestion was performed using RNase-free DNase (Qiagen) to remove contaminating genomic DNA. RNA samples were quantified using a Nanodrop Spectrophotometer (Thermo Scientific) and their quality was evaluated using a 2100 Bioanalyzer (Agilent Technologies). All RNA samples had an RNA integrity number ≥ 9.6. RNA samples were processed for microarray using the GeneChip^®^ 3’ IVT Express Kit (Affymetrix) according to the manufacturer’s instructions as previously^48^. Total RNA (100 ng) was used as input and quality checks were performed using the Bioanalyzer at all stages. RNA samples were processed in batches of 12 and batch mixing was used at every stage to avoid creating experimental bias. Hybridisation of RNA to chips and scanning of arrays was performed by the Medical Research Council’s Clinical Sciences Centre (CSC) Genomics Laboratory, Hammersmith Hospital, London, UK as previously. RNA was hybridised to GeneChip Chicken Genome Array chips (Affymetrix; containing comprehensive coverage of 32,773 transcripts corresponding to over 28,000 chicken genes) in a GeneChip Hybridization Oven (Affymetrix), the chips were stained and washed on a GeneChip Fluidics Station 450 (Affymetrix), and the arrays were scanned in a GeneChip Scanner 3000 7G with autoloader (Affymetrix).

### Microarray data analysis

A one-way ANOVA adjusted with the Benjamini–Hochberg multiple-testing correction (false discovery rate (FDR) of *P*<0.05) was performed with Partek Genomics Suite (v6.6, Partek) across all samples as previously^48,49^. Comparisons were conducted between infected cells versus mock-treated cells and between infected cells with the KO viruses versus CEF infected with the parental FP9 strain. The analysis cut off criteria were fold change ≥ ±1.5 and *P*-value ≤ 0.05. The Affymetrix chicken genome arrays contain probe sets for detecting transcripts from 17 avian viruses, including FWPV, allowing confirmation of viral infection. Visualisation of gene expression data was conducted with GeneSpring GX (v.13.1, Agilent Technologies) and GraphPad Prism (v.6.0). Original microarray data produced in this study have been deposited according to the MIAME guidelines in the public database ArrayExpress (http://www.ebi.ac.uk/microarray-as/ae/) (Acc. No: E-MTAB-7276). The catalogue of 337 ISGs was created by applying a fold change>3 compared to mock and false discovery rate (FDR)<0.05 on previously published microarray data [^25^; ArrayExpress accession: E-MTAB-3711 and Supplementary Table S1].

### Quantitative real-time RT PCR

Quantitative real-time RT PCR was performed on RNA samples using a two-step procedure as previously^50^. RNA was first reverse-transcribed into cDNA using the QuantiTect Reverse Transcription Kit (Qiagen) according to manufacturer’s instructions. qPCR was then conducted on the cDNA in a 384-well plate with an ABI-7900HT Fast qPCR system (Applied Biosystems). Mesa Green qPCR MasterMix (Eurogentec) was added to the cDNA (5 μl for every 2 μl of cDNA). The following amplification conditions were used: 95°C for 5 min; 40 cycles of 95°C for 15 sec, 57°C for 20 sec, and 72°C for 20 sec; 95°C for 15 sec; 60°C for 15 sec; and 95°C for 15 sec. Primer sequences for genes that were used in the study are shown in S6. The output Ct values and dissociation curves were analysed using SDS v2.3 and RQ Manager v1.2 (Applied Biosystems). Gene expression data were normalized against the housekeeping gene GAPDH, and compared with the mock controls using the comparative C_T_ method [also referred to as the 2^−ΔΔCT^ method^51^]. All samples were loaded in triplicate.

### Confocal and widefield fluorescence microscopy

Immunofluorescence labelling was carried out using CEF seeded at 2.5 × 10^5^ cells/well on coverslips, incubated in 6 well plates at 37°C, 5% CO_2_ and infected with 0.5 - 1 pfu of virus for 24 h. Medium was aspirated, cells washed 3x with PBS and fixed with 4% paraformaldehyde in PBS at room temperature (RT). Coverslips were washed 3x in PBS and cells permeabilised (0.5% Triton X-100 in PBS) at RT for 10 min with shaking. Following a further 3x PBS washes, cells were blocked (0.5% bovine serum albumen in PBS) for 1 h at RT with shaking. Primary antibodies were applied at 1:200 in blocking solution for 1 h at RT with shaking followed by 5× 5 min PBS washes at RT. Secondary Alexa dye conjugated antibodies (Molecular Probes) were applied at 1:200 in blocking solution for 1 h at RT in the dark, followed by 5× 5 min PBS washes. For double-labelling experiments, a second round of incubation with primary antibody, washing and secondary antibody was carried out. To label DNA within cells, coverslips were incubated with 1:5000 TOPRO3 (Molecular Probes) for 10 min at RT, followed by 3x PBS washes. Coverslips were dipped in SuperQ water briefly, drained and mounted on Vectorshield mounting media (Vector labs, Burlingame, CA). Coverslips mounted with hard-set mounting media were allowed to set at 4°C overnight; all other coverslips were sealed with nail varnish. Confocal microscopy was performed using a Leica TCS NT confocal microscope (Leica, Heidelberg, Germany).

For widefield fluorescence microscopy, CEF were washed 2x with PBS at RT and fixed with 10% buffered formaldehyde. The images were acquired on Evos fluorescence microscope (Evos FL Imaging System, Thermo Fisher Scientific, USA).

### Virion fractionation

Purified GFP184 particles were incubated in a reaction mixture containing 50 mM Tris-HCl, pH 7.5 and 1% (vol/vol) NP-40, with or without 50 mM DTT for 1 h at 37°C. The insoluble and soluble materials were separated by centrifugation at 20,000x *g* for 30 min at 4°C.

### Western blotting

Proteins for western blots were harvested from CEF and DF-1 cells. Cell pellets were lysed with CelLytic-M solution (Sigma-Aldrich) and the supernatant collected by centrifugation at 15,000 xg, 15 min, prior to protease inhibitor (Roche) addition. To every 20 μl of sample, 5 μl of 4X loading buffer (Bio-Rad) was added, and the samples were heated at 60°C for 5 min. They were then separated on a 12% sodium dodecylsulfate polyacrylamide gel, alongside a protein ladder (Chameleon Dual Colour Standards, LICOR). Samples (20 μg) were loaded for each well, and the gel was run at 150V for 2 h. Protein samples fractionated by SDS-PAGE were electroblotted onto a nitrocellulose membrane (Hybond-ECL; Amersham Biosciences) by following standard protocols. After transfer, the membranes, blocked overnight in 5% nonfat dry milk in PBSa buffer, were incubated for 2 h at room temperature with anti-GFP (Sigma-Aldrich), mouse monoclonal anti-FLAG (M2) (Sigma-Aldrich), rabbit monoclonal a-tubulin (Cell signalling Technology), mouse polyclonal a-chicken Mx1 (AbMart), and mouse monoclonal a-FPV168 (GB9), a-FPV140 (DF6) and a-FPV191 (DE9) antibodies^27,45^ [in PBS + 0.1% Tween-20 (Sigma) + 2% non-fat dried skimmed milk] at a dilution of 1:(1000-5000), followed by washing with PBS five times for 5 min. Secondary antibody [goat anti-rabbit horseradish peroxidase conjugated or goat anti-rabbit or donkey anti-mouse secondary antibodies (LICOR)] was diluted in PBS + 0.1% Tween-20 and added to the blot. Incubation was allowed to proceed for 1 h followed by washing with PBS five times for 5 min each. Labelled proteins were detected by incubation with either the ECL detection reagent (Amersham Biosciences), and exposure to hyperfilm ECL chemiluminescence film (Amersham Biosciences) or scanned with the Odyssey Imaging system (LICOR).

### Transfection of cells with polyI:C and assay of luciferase reporters

Chicken fibroblast DF-1 cells were transfected in 12-well plates with either chicken Mx1 or IFN-β promoter reporters [100 ng^52^], the constitutive reporter plasmid pJATlacZ [100 ng^53^] or co-transfected with plasmids driving the overexpression of FPV184 or FPV012 or VV A19 or the control empty vector. Following recovery for 24 h, cells were either left untreated or treated with 1000 IU/ml recombinant IFN-α treatment and incubated for 6 h or transfected overnight with polyI:C (10 μg/ml) using Polyfect (Qiagen). Luciferase assays were carried out, and data were normalized using β-galactosidase measurements and expressed as fold induction over control.

### Correlative super-resolution light and electron microscopy (CSRLEM)

GFP184 viruses were pelleted through 36% sucrose and then band purified on a 25 to 40 % sucrose gradient as described previously^54^. Virions were diluted in 20 μl 1 mM Tris pH 9, placed in the centre of the clean coverslips for 30 min and bound virus was fixed with 4% EM-grade formaldehyde (TAAB). A small asymmetric scratch was made in the middle of the coverslip using a diamond scorer, to enable localisation of the super-resolution imaging region of interest within the resin block for trimming, targeting and subsequently in sections during electron imaging. Sample was permeabilized for 30 min with 1% Triton X-100 in PBS and blocked with PBS containing 5% BSA (Sigma), 1% FCS for 30 min. Sample was immunostained overnight with anti-GFP nanobody (Chromotek) conjugated in-house to AlexaFluor647-NHS (Invitrogen) in PBS containing 5% BSA at 4°C. Coverslips were washed 3 times with PBS and mounted on a microscope slide with parafilm gasket in 1% (v/v) β-mercaptoethanol (Sigma), 150 mM Tris, 1% glucose, 1% glycerol, 10 mM NaCl, pH 8 with 0.25 mg/ml glucose-oxidase and 20 μg/ml catalase.

Super-resolution microscopy was performed on an Elyra PS.1 inverted microscope (Zeiss) using an alpha Plan-Apochromat 100× /1.46 NA oil DIC M27 objective. STORM^55^ images were acquired with a 1.6x tube lens on an iXon 897 EMCCD camera (Andor) with 20ms exposure time with 642nm excitation at 100% laser power and a 655nm LP filter. Fluorophore activation was dynamically controlled with a 405 nm laser at 0-2% laser power. Images were processed in Fiji^56^ using ThunderSTORM^57^. Localizations were fitted with a maximum-likelihood estimator, lateral drift corrected by cross-correlation, localizations <20nm apart within ≤1 frames merged, and images rendered using a Gaussian profile.

After super-resolution imaging of the region of interest, a series of phase contrast images using objectives with 10x, 20x, 40x magnification and fluorescence images using an objective with 100x magnification were taken to map the region of interest and its localisation in relation to the scratch. Coverslips were washed twice in PBS and fixed with 1.5% glutaraldehyde and 2% EM-grade paraformaldehyde (TAAB) in 0.1M sodium cacodylate for 45 min at room temperature. Samples were treated with reduced osmium and tannic acid, dehydrated through an ethanol series and embedded in epon resin, as previously described^58^. After resin polymerisation, the coverslip was removed using liquid nitrogen, revealing the positive pattern of the coverslip-scratch in the surface of the resin block. Using the scratch mark as reference and viral clusters from phase images as fiducials, the region of interest that had been imaged for super-resolution imaging, was identified and targeted for serial sectioning. Sections were collected on formvar coated slot grids, stained with lead citrate and imaged using a transmission electron microscope (Tecnai T12, Thermo Fisher Scientific) equipped with a charge-coupled device camera (SIS Morada; Olympus). EM and STORM images were registered using NanoJ^59^. It should be noted that preparation of samples for EM after super-resolution microscopy may move or remove individual virus particles, which mandates careful registration and manual overlay of the images. However, the orientation and structure of the fluorescence signal are clearly reminiscent of the lateral body structure observed in EM.

### Phylogenetic analysis

The amino acid sequences of FPV184 orthologues from each genera of chordopoxviruses were subjected to multiple alignments (Fig. 5a & Supplementary Fig. S5) using CLC workbench 7 (CLC Bio, Qiagen, Aarhus, Denmark). Protein sequence accession numbers for the indicated viruses are as follows: fowlpox virus (FPV184, NP_039147), canarypox Virus (CNPV258, NP_955281), VACV (A19, P68714), myxoma Virus (m109L, AQT34599), deerpox virus (DpV83gp120, YP_227495), sheeppox virus (SPPV_106, NP_659683), swinepox Virus (SPV108, NP_570268), Yaba monkey tumor virus (111L, NP_938366), molluscum contagiosum virus (MC124, AQY16697), orf virus (ORF096, NP_957873), crocodilepox virus (P4b, YP_784314).

### Statistical analysis

To determine the significance of differences between experimental groups, one- or two-way ANOVA analyses were performed using the fold change scores with a Tukey’s or Dunnett’s multiple comparisons test depending on the application^60^. *P*-values were set at 0.05 (*P*≤0.05) unless indicated otherwise. Error bars represent the standard error of the mean (SE). The correlation of expression values between microarray analysis and qRT-PCR was statistically assessed by calculation of Pearson’s correlation coefficient using the built-in function of GraphPad Prism (v.6.0).

## Data availability

Microarray data are available at the public database ArrayExpress (http://www.ebi.ac.uk/microarray-as/ae/) (Accession Number: E-MTAB-7276). All other data supporting the findings of this study are available from the corresponding author on request.

## Acknowledgements

Our thanks go to Steve Goodbourn, Jack Hankinson, Laurence Game, Nathalie Lambie and Ivan Andrew for their technical assistance. This research was undertaken with the financial support of the Biotechnology and Biological Sciences Research Council (BBSRC) (http://www.bbsrc.ac.uk) *via* grants BB/H005323/1 (“Correlation of immunogenicity with microarray analysis of vector mutants to improve live recombinant poxvirus vaccines in poultry”), BB/E009956/1 (“Viral & host immunomodulators in improved Fowlpox virus recombinant vector vaccines for use in poultry against highly pathogenic Avian Influenza H5N1”) and BB/K002465/1 (Strategic LoLa; “Developing Rapid Responses to Emerging Virus Infections of Poultry (DRREVIP)”. The CSRLEM work was funded by MRC Programme grant (MC_UU_00012/7; to JM) and the European Research Council (649101, UbiProPox; to JM). SRB is a Sir Henry Wellcome Post-doctoral Fellow (WT106080/Z/14/Z). DA is a Marie Skłodowska-Curie fellow funded by the European Union (750673). JJB is supported by MRC funding to the MRC LMCB University Unit at UCL, award code MC_U12266B. ESG is also supported by funding from the Wellcome Trust new investigator award (104771/Z/14/Z).

## Author contributions

Conceived and designed the experiments: ESG, SML, SRB, DA, JM, MAS. Performed the experiments: ESG, SML, SRB, DA, JJB, RCR. Analysed the data: ESG, SML, SRB, DA, JJB, RCR, JM, MAS. Wrote the paper: ESG, SML, SRB, DA, JJB, JM, MAS.

## Additional Information

The authors declare no competing financial interests.

## References

1 McFadden, G. Poxvirus tropism. Nat Rev Microbiol 3, 201–213 (2005).

2 Hruby, D. E., Lynn, D. L. & Kates, J. R. Vaccinia virus replication requires active participation of the host cell transcriptional apparatus. Proc Natl Acad Sci U S A 76, 1887–1890 (1979).

3 Hruby, D. E., Guarino, L. A. & Kates, J. R. Vaccinia virus replication. I. Requirement for the host-cell nucleus. J Virol 29, 705–715 (1979).

4 Pennington, T. H. & Follett, E. A. Vaccinia virus replication in enucleate BSC-1 cells: particle production and synthesis of viral DNA and proteins. J Virol 13, 488–493 (1974).

5 Prescott, D. M., Kates, J. & Kirkpatrick, J. B. Replication of vaccinia virus DNA in enucleated L-cells. J Mol Biol 59, 505–508 (1971).

6 Prescott, D. M., Myerson, D. & Wallace, J. Enucleation of mammalian cells with cytochalasin B. Exp Cell Res 71, 480–485 (1972).

7 Sen, G. C. Viruses and interferons. Annu Rev Microbiol 55, 255–281 (2001).

8 Katze, M. G., He, Y. & Gale, M., Jr. Viruses and interferon: a fight for supremacy. Nat Rev Immunol 2, 675–687 (2002).

9 Seet, B. T. et al. Poxviruses and immune evasion. Annu Rev Immunol 21, 377–423 (2003).

10 Bidgood, S. R. & Mercer, J. Cloak and Dagger: alternative immune evasion and modulation strategies of poxviruses. Viruses 7, 4800–4825 (2015).

11 Bhattacharya, S. et al. Anti-tumorigenic effects of Type 1 interferon are subdued by integrated stress responses. Oncogene 32, 4214–4221 (2013).

12 Smith, G. L. et al. Vaccinia virus immune evasion: mechanisms, virulence and immunogenicity. J Gen Virol 94, 2367–2392 (2013).

13 Paludan, S. R., Bowie, A. G., Horan, K. A. & Fitzgerald, K. A. Recognition of herpesviruses by the innate immune system. Nat Rev Immunol 11, 143–154 (2011).

14 Xing, J. et al. Herpes simplex virus 1-encoded tegument protein VP16 abrogates the production of beta interferon (IFN) by inhibiting NF-kappaB activation and blocking IFN regulatory factor 3 to recruit its coactivator CBP. J Virol 87, 9788–9801 (2013).

15 Harle, P., Sainz, B., Jr., Carr, D. J. & Halford, W. P. The immediate-early protein, ICP0, is essential for the resistance of herpes simplex virus to interferon-alpha/beta. Virology 293, 295–304 (2002).

16 Jiang, Z., Su, C. & Zheng, C. Herpes simplex virus 1 tegument protein UL41 counteracts IFIT3 antiviral innate immunity. J Virol 90, 11056–11061 (2016).

17 Schmidt, F. I. et al. Vaccinia virus entry is followed by core activation and proteasome-mediated release of the immunomodulatory effector VH1 from lateral bodies. Cell Rep 4, 464–476 (2013).

18 Najarro, P., Traktman, P. & Lewis, J. A. Vaccinia virus blocks gamma interferon signal transduction: viral VH1 phosphatase reverses Stat1 activation. J Virol 75, 3185–3196 (2001).

19 Giotis, E. S. & Skinner, M. A. Spotlight on avian pathology: fowlpox virus. Avian Pathol 48, 87–90 (2019).

20 Laidlaw, S. M. et al. Genetic screen of a mutant poxvirus library identifies an ankyrin repeat protein involved in blocking induction of avian type I interferon. J Virol 87, 5041–5052 (2013).

21 Laidlaw, S. M. & Skinner, M. A. Comparison of the genome sequence of FP9, an attenuated, tissue culture-adapted European strain of Fowlpox virus, with those of virulent American and European viruses. J Gen Virol 85, 305–322 (2004).

22 Skinner, M. A., Laidlaw, S. M., Eldaghayes, I., Kaiser, P. & Cottingham, M. G. Fowlpox virus as a recombinant vaccine vector for use in mammals and poultry. Expert Rev Vaccines 4, 63–76 (2005).

23 Laidlaw, S. M. & Skinner, M. A. Construction of deletion-knockout mutant fowlpox virus (FWPV). Bio Protoc 4(2014).

24 Buttigieg, K. et al. Genetic screen of a library of chimeric poxviruses identifies an ankyrin repeat protein involved in resistance to the avian type I interferon response. J Virol 87, 5028–5040 (2013).

25 Giotis, E. S. et al. Chicken interferome: avian interferon-stimulated genes identified by microarray and RNA-seq of primary chick embryo fibroblasts treated with a chicken type I interferon (IFN-alpha). Vet Res 47, 75 (2016).

26 Ichihashi, Y. & Oie, M. The activation of vaccinia virus infectivity by the transfer of phosphatidylserine from the plasma membrane. Virology 130, 306–317 (1983).

27 Boulanger, D. et al. Identification and characterization of three immunodominant structural proteins of fowlpox virus. J Virol 76, 9844–9855 (2002).

28 Szajner, P., Weisberg, A. S., Wolffe, E. J. & Moss, B. Vaccinia virus A30L protein is required for association of viral membranes with dense viroplasm to form immature virions. J Virol 75, 5752–5761 (2001).

29 Satheshkumar, P. S., Olano, L. R., Hammer, C. H., Zhao, M. & Moss, B. Interactions of the vaccinia virus A19 protein. J Virol 87, 10710–10720 (2013).

30 Satheshkumar, P. S., Weisberg, A. S. & Moss, B. Vaccinia virus A19 protein participates in the transformation of spherical immature particles to barrel-shaped infectious virions. J Virol 87, 10700–10709 (2013).

31 Brown, M. et al. Dendritic cells infected with recombinant fowlpox virus vectors are potent and long-acting stimulators of transgene-specific class I restricted T lymphocyte activity. Gene Ther 7, 1680–1689 (2000).

32 Miller, J., McLachlan, A. D. & Klug, A. Repetitive zinc-binding domains in the protein transcription factor IIIA from Xenopus oocytes. EMBO J 4, 1609–1614 (1985).

33 Orkin, S. H. GATA-binding transcription factors in hematopoietic cells. Blood 80, 575–581 (1992).

34 Lentjes, M. H. et al. The emerging role of GATA transcription factors in development and disease. Expert Rev Mol Med 18, e3 (2016).

35 Hu, X., Carroll, L. J., Wolffe, E. J. & Moss, B. De novo synthesis of the early transcription factor 70-kilodalton subunit is required for morphogenesis of vaccinia virions. J Virol 70, 7669–7677 (1996).

36 Hu, X., Wolffe, E. J., Weisberg, A. S., Carroll, L. J. & Moss, B. Repression of the A8L gene, encoding the early transcription factor 82-kilodalton subunit, inhibits morphogenesis of vaccinia virions. J Virol 72, 104–112 (1998).

37 Smiley, J. R. Herpes simplex virus virion host shutoff protein: immune evasion mediated by a viral RNase? J Virol 78, 1063–1068 (2004).

38 Stuart, J. H., Sumner, R. P., Lu, Y., Snowden, J. S. & Smith, G. L. Vaccinia Virus Protein C6 Inhibits Type I IFN Signalling in the Nucleus and Binds to the Transactivation Domain of STAT2. PLoS Pathog 12, e1005955 (2016).

39 Senkevich, T. G., Koonin, E. V. & Moss, B. Vaccinia virus F16 protein, a predicted catalytically inactive member of the prokaryotic serine recombinase superfamily, is targeted to nucleoli. Virology 417, 334–342 (2011).

40 Oh, J. & Broyles, S. S. Host cell nuclear proteins are recruited to cytoplasmic vaccinia virus replication complexes. J Virol 79, 12852–12860 (2005).

41 Rosales, R., Sutter, G. & Moss, B. A cellular factor is required for transcription of vaccinia viral intermediate-stage genes. Proc Natl Acad Sci U S A 91, 3794–3798 (1994).

42 Camus-Bouclainville, C. et al. A virulence factor of myxoma virus colocalizes with NF-kappaB in the nucleus and interferes with inflammation. J Virol 78, 2510–2516 (2004).

43 Rodriguez, J. A., Span, S. W., Kruyt, F. A. & Giaccone, G. Subcellular localization of CrmA: identification of a novel leucine-rich nuclear export signal conserved in anti-apoptotic serpins. Biochem J 373, 251–259 (2003).

44 Laidlaw, S. M. et al. Fowlpox virus encodes nonessential homologs of cellular alpha-SNAP, PC-1, and an orphan human homolog of a secreted nematode protein. J Virol 72, 6742–6751 (1998).

45 Boulanger, D., Green, P., Smith, T., Czerny, C. P. & Skinner, M. A. The 131-amino-acid repeat region of the essential 39-kilodalton core protein of fowlpox virus FP9, equivalent to vaccinia virus A4L protein, is nonessential and highly immunogenic. J Virol 72, 170–179 (1998).

46 Irwin, C. R. & Evans, D. H. Modulation of the myxoma virus plaque phenotype by vaccinia virus protein F11. J Virol 86, 7167–7179 (2012).

47 Falkner, F. G. & Moss, B. Transient dominant selection of recombinant vaccinia viruses. J Virol 64, 3108–3111 (1990).

48 Giotis, E. S. et al. Constitutively elevated levels of SOCS1 suppress innate responses in DF-1 immortalised chicken fibroblast cells. Sci Rep 7, 17485 (2017).

49 Giotis, E. S. et al. Entry of the bat influenza H17N10 virus into mammalian cells is enabled by the MHC class II HLA-DR receptor. Nat Microbiol (2019).

50 Giotis, E. S. et al. Transcriptomic Profiling of virus-host cell interactions following chicken anaemia virus (CAV) infection in an in vivo model. PLoS One 10, e0134866 (2015).

51 Pfaffl, M. W. A new mathematical model for relative quantification in real-time RT-PCR. Nucleic Acids Res 29, e45 (2001).

52 Childs, K. et al. mda-5, but not RIG-I, is a common target for paramyxovirus V proteins. Virology 359, 190–200 (2007).

53 Masson, N., Ellis, M., Goodbourn, S. & Lee, K. A. Cyclic AMP response element-binding protein and the catalytic subunit of protein kinase A are present in F9 embryonal carcinoma cells but are unable to activate the somatostatin promoter. Mol Cell Biol 12, 1096–1106 (1992).

54 Yakimovich, A. et al. Inhibition of poxvirus gene expression and genome replication by bisbenzimide derivatives. J Virol 91(2017).

55 Heilemann, M. et al. Subdiffraction-resolution fluorescence imaging with conventional fluorescent probes. Angew Chem Int Ed Engl 47, 6172–6176 (2008).

56 Schindelin, J. et al. Fiji: an open-source platform for biological-image analysis. Nat Methods 9, 676–682 (2012).

57 Ovesny, M., Krizek, P., Borkovec, J., Svindrych, Z. & Hagen, G. M. ThunderSTORM: a comprehensive ImageJ plug-in for PALM and STORM data analysis and super-resolution imaging. Bioinformatics 30, 2389–2390 (2014).

58 Agrotis, A., Pengo, N., Burden, J. J. & Ketteler, R. Redundancy of human ATG4 protease isoforms in autophagy and LC3/GABARAP processing revealed in cells. Autophagy, 1–22 (2019).

59 Laine, R. F. et al. NanoJ: a high-performance open-source super-resolution microscopy toolbox. Journal of Physics D: Applied Physics 52(2019).

60 Lee, S. & Lee, D. K. What is the proper way to apply the multiple comparison test? Korean J Anesthesiol 71, 353–360 (2018).

